## Supplementary figures and images for "Host JAK-STAT activity is a target of parasitoid wasp virulence strategies"

### Fig S1

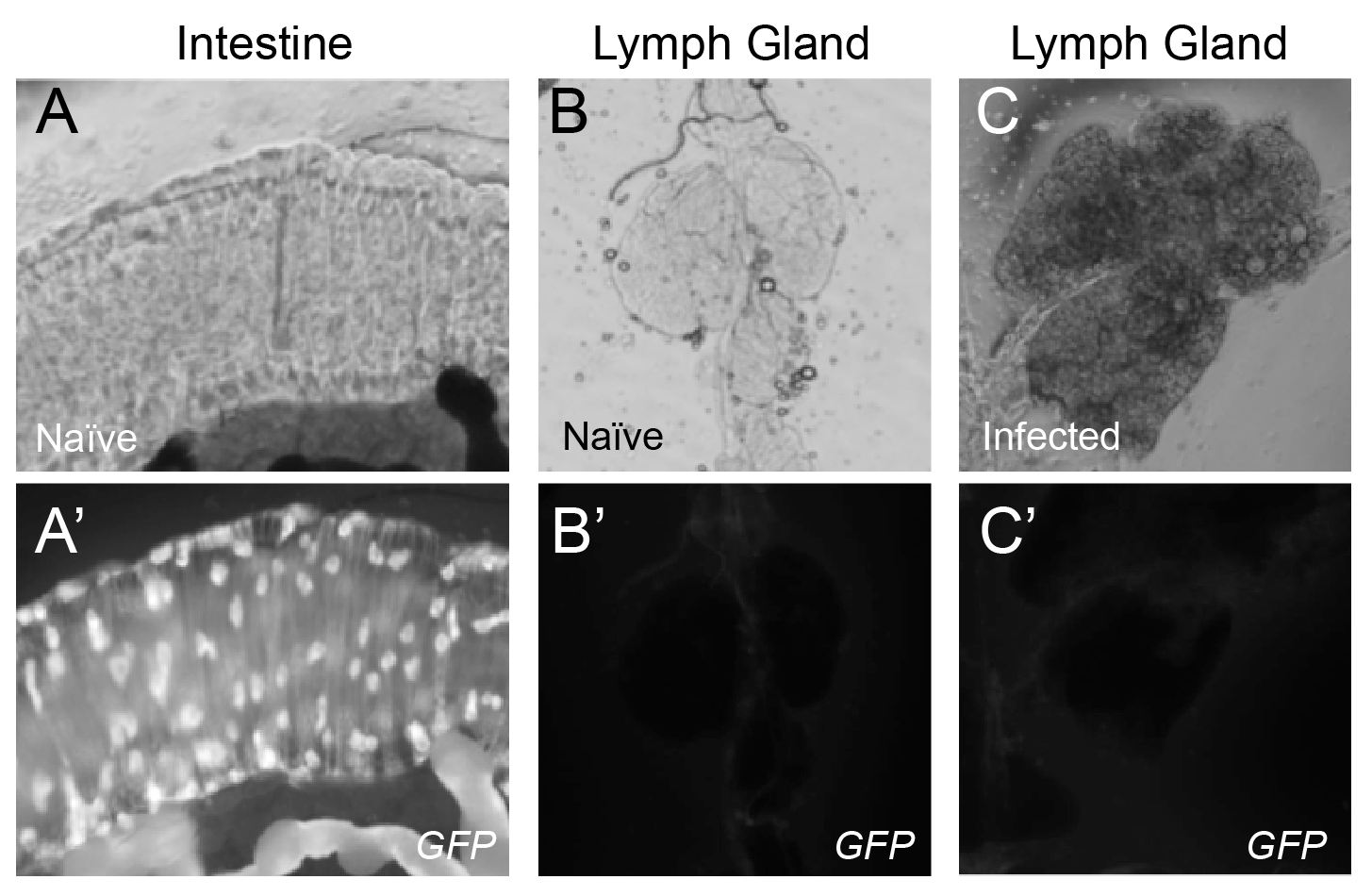

### Fig S2

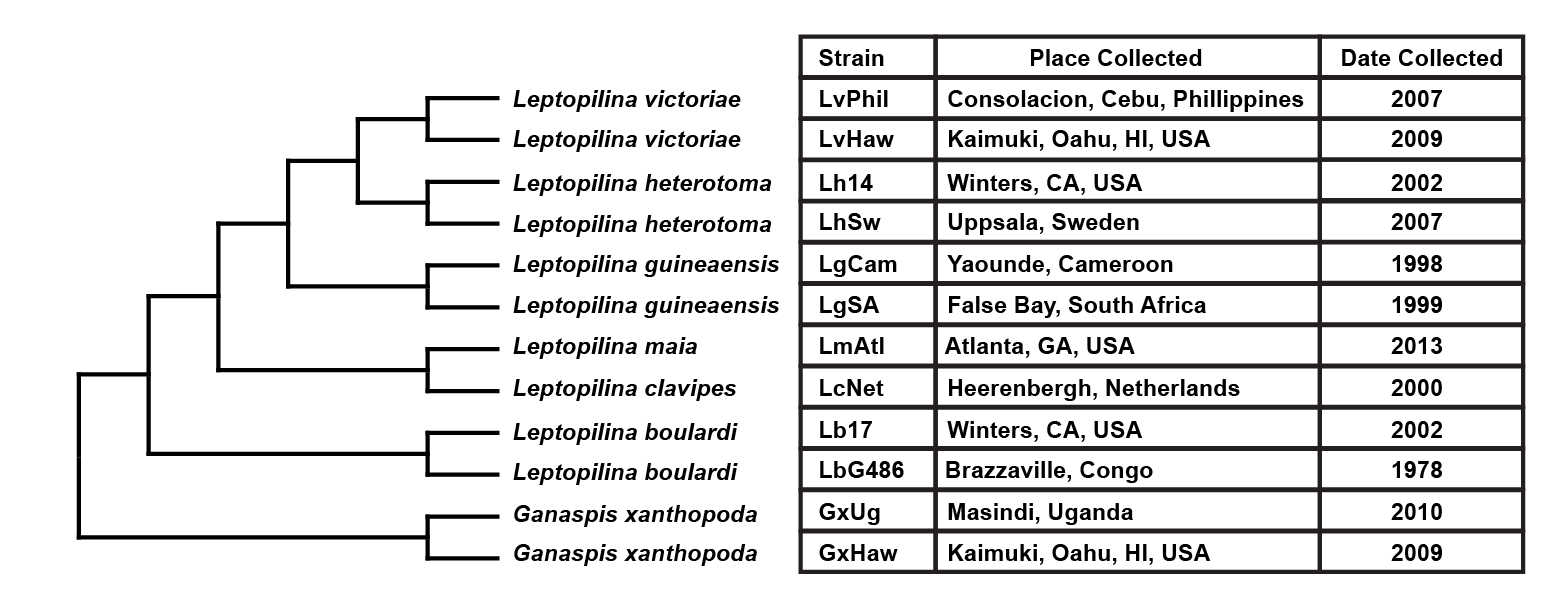

### Fig S3

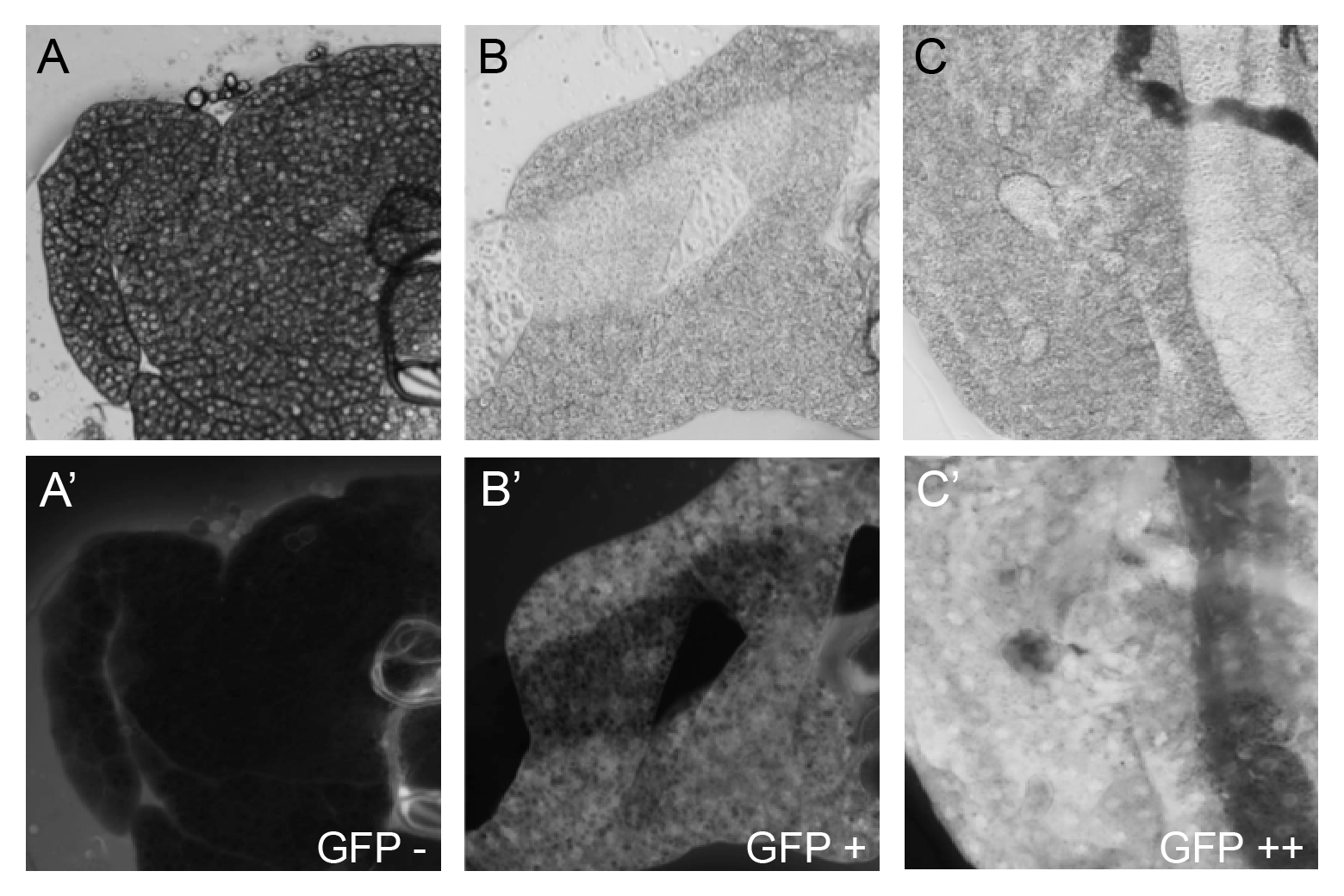

### Fig S4

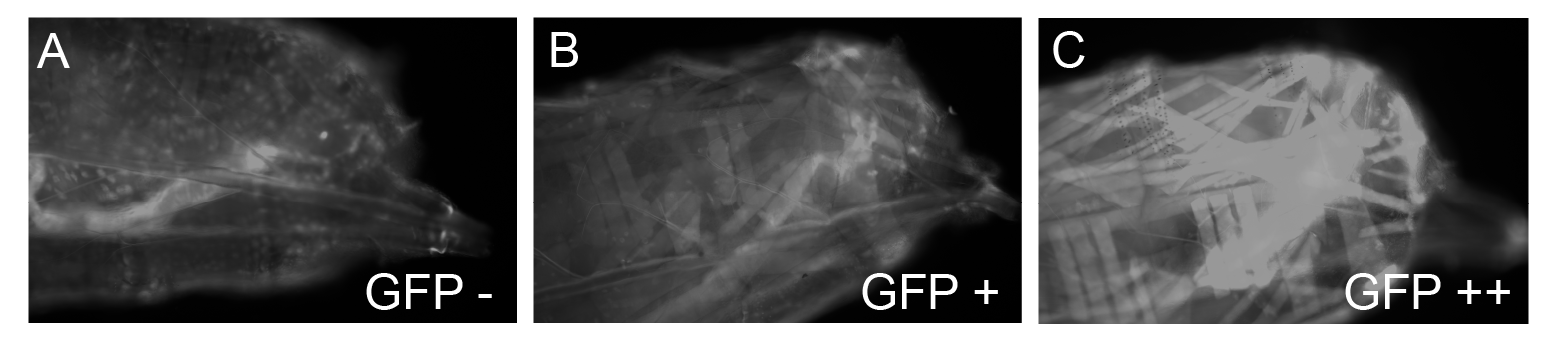
