## Supplemental Table 1 for "Host JAK-STAT activity is a target of parasitoid wasp virulence strategies"

**Supplemental Table 1.** Species and strains names, and COI sequence accession numbers for all parasitoids used in this study.

| Species | Strain | Accession # |
| --- | --- | --- |
| *Leptopilina victoriae* | LvPhil | JQ808447 |
| *Leptopilina victoriae* | LvHaw | JQ808446 |
| *Leptopilina heterotoma* | Lh14 | JQ808444 |
| *Leptopilina heterotoma* | LhSw | JQ808445 |
| *Leptopilina guineaensis* | LgCam | JQ808442 |
| *Leptopilina guineaensis* | LgSA | JQ808443 |
| *Leptopilina maia* | LmAtl | JQ808440 |
| *Leptopilina clavipes* | LcNet | JQ808441 |
| *Leptopilina boulardi* | Lb17 | JQ808436 |
| *Leptopilina boulardi* | LbG486 | JQ808438 |
| *Ganaspis xanthopoda* | GxUg | JQ808434 |
| *Ganaspis xanthopoda* | GxHaw | JQ808433 |

**Supplemental Figure Legends**

**Supplemental Figure 1.** Brightfield (A-C) and fluorescence (A’-C’) images of *10xSTAT92E-GFP* larvae. Strong fluorescence is seen in gut cells (A,A’). No fluorescence in seen in lymph gland dissected from naïve (B,B’) or LcNet infected (C,C’) larvae.

**Supplemental Figure 2.** Phylogenetic tree of parasitoid species and strains used in this study. Phylogeny was constructed using COI sequences given in Supplemental Table 1. The place and date of collection is also given for each strain.

**Supplemental Figure 3.** Brightfield (A-C) and fluorescence (A’-C’) images of fat bodies dissected from *10xSTAT92E-GFP* larvae. Images are representative of the expression categories used in Table 1. A’ is representative of ‘-‘, B’ is representative of ‘+’ and C’ is representative of ‘++’.

**Supplemental Figure 4.** Fluorescent images of body wall muscle from *10xSTAT92E-GFP* larvae. Images are representative of the expression categories used in Table 2. A is representative of ‘-‘, B is representative of ‘+’ and C is representative of ‘++’. All images were taken at the posterior end.
